# Zebrafish *her3* knockout impacts developmental and cancer-related gene signatures

**DOI:** 10.1101/2022.02.25.482038

**Authors:** Matthew R. Kent, Delia Calderon, Katherine M. Silvius, Collette A. LaVigne, Matthew V. Cannon, Genevieve C. Kendall

**Affiliations:** Center for Childhood Cancer & Blood Diseases, The Abigail Wexner Research Institute, Nationwide Children’s Hospital, Columbus, OH 43205, USA; Molecular, Cellular, and Developmental Biology Ph.D. Program, The Ohio State University, Columbus, OH 43210, USA; Department of Molecular Biology, UT Southwestern Medical Center, Dallas, TX 75390, USA; Department of Pediatrics, The Ohio State University College of Medicine, Columbus, OH 43205, USA

**Keywords:** her3/HES3, functional genomics, zebrafish, CRISPR/Cas9 knockout, neural development, transcriptomics

## Abstract

*HES3* is a basic helix-loop-helix transcription factor that regulates neural stem cell renewal during development. *HES3* overexpression is predictive of reduced overall survival in patients with fusion-positive rhabdomyosarcoma, a pediatric cancer that resembles immature and undifferentiated skeletal muscle. However, the mechanisms of *HES3* cooperation in fusion-positive rhabdomyosarcoma are unclear and are likely related to *her3*/*HES3’s* role in neurogenesis. To investigate *HES3’s* function during development, we generated a zebrafish CRISPR/Cas9 knockout of *her3*, the zebrafish ortholog of *HES3*. Loss of *her3* is not embryonic lethal and adults exhibit expected Mendelian ratios. Embryonic *her3* zebrafish mutants are significantly smaller than wildtype and a subset present with lens defects as adults. Transcriptomic analysis of *her3* mutant embryos indicates that genes involved in organ development, such as *pctp and grinab*, are significantly downregulated. Further, differentially expressed genes in *her3* knockout embryos are enriched for HOX and SOX10 motifs. Several cancer-related gene pathways are impacted, including the inhibition of matrix metalloproteinases. Altogether, this new model is a powerful system to study *her3/HES3*-mediated neural development and its misappropriation in cancer contexts.

**Summary Statement:** Here, we generate and characterize a zebrafish *her3*/*HES3* knockout to elucidate the functional role of *her3*/*HES3*, a transcriptional repressor, in neural development and tumorigenic processes.

## Introduction

*HES3*, and the zebrafish ortholog *her3*, are part of the *HES* gene family, a group of basic helix-loop-helix transcriptional repressors (Sasai et al., 1992). There are seven *HES* family genes that are conserved between humans and mice, in addition to *HES4*, which contains no mouse ortholog. In zebrafish, there are 13 *her* genes that are orthologs to the mammalian *HES* genes: *her6* is orthologous to *HES1*; *her3* is orthologous to *HES3*; *her9* to *HES4*; *her2, her4*.*1, her12*, and *her15* to *HES5*; *her8a* and *her13* to *HES6*; and *her1, her5, her7*, and *her11* to *HES7* (Davis and Turner, 2001, El Yakoubi et al., 2012, Sieger et al., 2004, Shankaran et al., 2007, Sieger et al., 2006, Gajewski and Voolstra, 2002). A subset of the *HES* family is known to regulate neurogenesis, such as *HES1, HES3*, and *HES5* (Ohtsuka et al., 1999, Hirata et al., 2001, Hatakeyama et al., 2004). These data originated from a *Hes1/Hes3* double knock-out mouse, in which neural progenitor cells in the midbrain and anterior hindbrain prematurely differentiated, preventing these areas from fully developing (Hirata et al., 2001). In mouse and rat neural stem cells, *HES3/her3* functions in a non-canonical Notch signaling pathway. This pathway is mediated through soluble Notch ligands which bind to Notch receptors, thus activating kinases to phosphorylate serine-727 on STAT3, and ultimately resulting in an upregulation of *HES3* and self-renewal of neural stem cells (Artavanis-Tsakonas et al., 1999, Androutsellis-Theotokis et al., 2006, Androutsellis-Theotokis et al., 2009, Kageyama et al., 2008, Hans et al., 2004, Bae et al., 2005).

Other *HES* family genes are responsible for segmentation of somites in mice, bilateral structures on either side of the neural tube that give rise to vertebrae, ribs, and skeletal muscle as examples (Dubrulle and Pourquie, 2004). Cyclic expression of *Hes1, Hes5*, and *Hes7* in mice, and *her1* and *her7* in zebrafish, are responsible for the segmentation of somites (Jouve et al., 2000, Dunwoodie et al., 2002, Bessho et al., 2001a, Bessho et al., 2001b, Muller et al., 1996, Holley et al., 2002, Oates and Ho, 2002). This is a tightly regulated developmental process, as cyclical expression of *HES7* is critical for somite segmentation, and in the absence of *HES7*, somites fuse (Bessho et al., 2001b). Interestingly, persistent activation of *HES7* also leads to somite fusion (Hirata et al., 2004). Another HES family member, *Hes6*, has been shown to regulate differentiation and fusion of myoblasts and its overexpression is predictive of poor prognosis in alveolar rhabdomyosarcoma (ARMS) (Gao et al., 2001, Malone et al., 2011, Wickramasinghe et al., 2013). These data suggest that HES genes have dual roles in mesodermal specification and myogenesis, but HES3, a neural transcription factor, has not been implicated in these processes.

Previously, we identified a role for HES3 in fusion-positive rhabdomyosarcoma, a pediatric cancer that resembles immature skeletal muscle and is characterized by an inability to terminally differentiate (Parham and Ellison, 2006). We found that injecting the human *PAX3-FOXO1* fusion, the defining oncogene in alveolar rhabdomyosarcoma (Barr et al., 1993, Barr, 2011), into zebrafish embryos significantly upregulated *her3* mRNA expression. Further, *her3* overexpression was unique to and dependent on *PAX3-FOXO1*. Our finding is translatable to the human disease, as *HES3* is overexpressed in fusion-positive alveolar rhabdomyosarcoma patient tumors, and this overexpression is predictive of reduced overall survival (Kendall et al., 2018). This suggests that *HES3* has functional consequences when expressed in fusion-positive ARMS. Given that other HES genes are implicated in myogenic and tumorigenic processes, we sought to understand how *her3*/*HES3* could be contributing to this disease.

Here, we make a series of *her3* CRISPR/Cas9-knockout mutant alleles in zebrafish to investigate *her3’s* role in developmental processes and probe how this could translate to its role during tumorigenesis. We show that loss of *her3* is not embryonic lethal and progeny exhibit the expected Mendelian ratios. In addition, *her3* knockout results in significant reduction of overall size in zebrafish embryos and lens degradation in a subset of fish as adults. Our RNA-seq analysis of differentially expressed genes of early zebrafish embryos suggests: 1) an enrichment of genes involved in the inhibition of matrix metalloproteinases; and 2) an enrichment for SOX10-regulated genes, a neural crest-specific transcription factor. These functions during neural development, stem cell maintenance, and extracellular matrix (ECM) homeostasis could be important for *her3/HES3*’s role as a cooperating gene in fusion-positive ARMS. Our new animal model is a powerful system to study these outstanding questions.

## Results

### Generating *her3* knockout mutations in zebrafish

To generate *her3* frameshift mutations in zebrafish, we used a CRISPR/Cas9 method adapted from the Talbot and Amacher, 2014 protocol (Talbot and Amacher, 2014). Wildtype WIK zebrafish embryos were injected at the 1-2 cell stage in the cell body with a *her3* gRNA and Cas9 protein (Figure 1A). The *her3* gRNA resulted in multiple potential mutations in P0 crispants, as determined by High Resolution Melt Analysis (HRMA) (Figure S1).). The injected embryos were allowed to develop to sexual maturity (∼ 3 months), and then were screened as potential founders by outcrossing to wildtype WIK and analyzing F1 offspring using HRMA. This strategy identified six potential founders from which one founder with the most diverse mutations was chosen to generate an F1 population (Figure S2). HRMA screening of the adult F1 heterozygous offspring revealed three potential mutations (Figure 1B). To determine the exact mutation, the region surrounding the gRNA binding site was A-tail cloned, sequenced, and analyzed via translation prediction (Figure 1C). The first mutation identified was a 4bp deletion, resulting in a frameshift after the 37^th^ amino acid, a 38^th^ altered amino acid, and a premature stop codon. The second mutation identified was a 7bp deletion resulting in a frameshift after the 38^th^ amino acid, 25 additional altered amino acids, and a premature stop codon. The third mutation identified was a 20bp insertion resulting in a frameshift after the 42^nd^ amino acid, 30 additional altered amino acids, and a premature stop codon. These are each referred throughout the text as: her3_38aa, her3_63aa, and her3_72aa. Each of these three mutations was a potential knockout, with the frameshift occurring in the middle of the DNA binding domain (Figure 1C).

**Figure 1:**
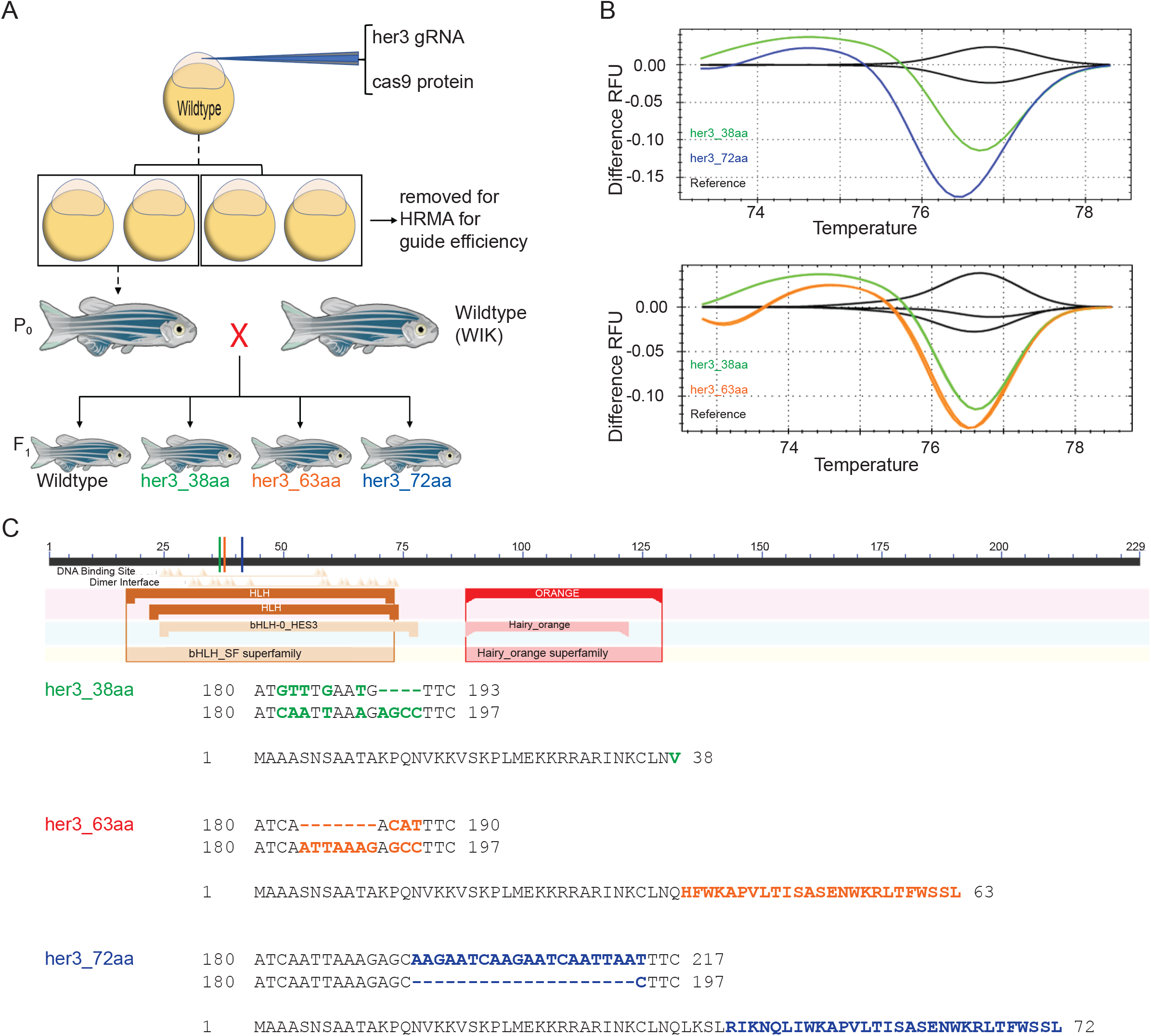
Generating *her3* mutations in zebrafish. (A) Schematic of strategy to generate a *her3* CRISPR/Cas9 knockout zebrafish. Zebrafish embryos were injected in the cell body at the 1-cell stage with both cas9 protein and *her3* gRNA. A subset of the injected clutch was set aside to determine gRNA efficiency. The remaining embryos were grown up to identify adult founders, which were outcrossed to wildtype WIK zebrafish to generate the F_1_ generation. (B) HRMA difference RFU graphs of F_1_ heterozygous zebrafish generated from three identified founders. Fin clips of adult F_1_ zebrafish were taken and HRMA was performed to identify potential mutations, with different melt curves indicating distinct DNA mutations. Wildtype reference fish are labeled black, her3_38aa fish are labeled green, her3_63aa fish are labeled orange, and her3_72aa fish are labeled blue. Each line represents an individual fish. (C) Schematic of full length *her3* protein and identified mutation sequences. To determine the mutation sequence, the *her3* gRNA target site from the three prospective mutant F_1_ fish was A-tail cloned into the pGEM T-Easy vector. Multiple colonies from each line were Sanger sequenced with an SP6 primer. The mutated sequence was then aligned with the wildtype reference sequence for *her3*. Colored lines in the schematic indicate where the predicted frameshift mutation occurs for each respective mutation, all of which are in the DNA binding domain. Below, the DNA sequence and amino acid sequence are shown for the three mutations, with mutant DNA sequence on top of the aligned wildtype sequence. For both the DNA sequence and the amino acid sequence, colored and bolded letters highlight the mutated sequence.

To verify that our mutants were indeed knockouts, we utilized an adapted reporter assay (Prykhozhij et al, 2017). We focused on the reporter assay because commercially available HES3 antibodies failed to recognize zebrafish her3 (data not shown). Zebrafish *her3* cDNA was amplified from either wildtype WIK or the three her3 homozygous mutant lines and cloned into the pCS2-MCS-P2A-sfGFP vector. RNA from one of these cloned constructs, as well as RNA from pCS2+TagRFPT as an injection control, was co-injected into the yolk of wildtype WIK embryos at the 1-4 cell stage (Figure 2A). At 24 hours post-fertilization (hpf), individual embryos were then imaged for GFP expression (*her3* construct) and RFP expression (control construct). Each of the three lines tested - her3_38aa, her3_63aa, and her3_72aa -were determined to be knockouts as shown by the significant decrease in GFP/RFP fluorescence ratio compared to the wildtype her3 RNA (Figures 2B-C). This ratio of GFP/RFP fluorescence was quantified for each her3 allele (Figure 2C) and was the basis for focusing on these three lines for further assessment.

**Figure 2:**
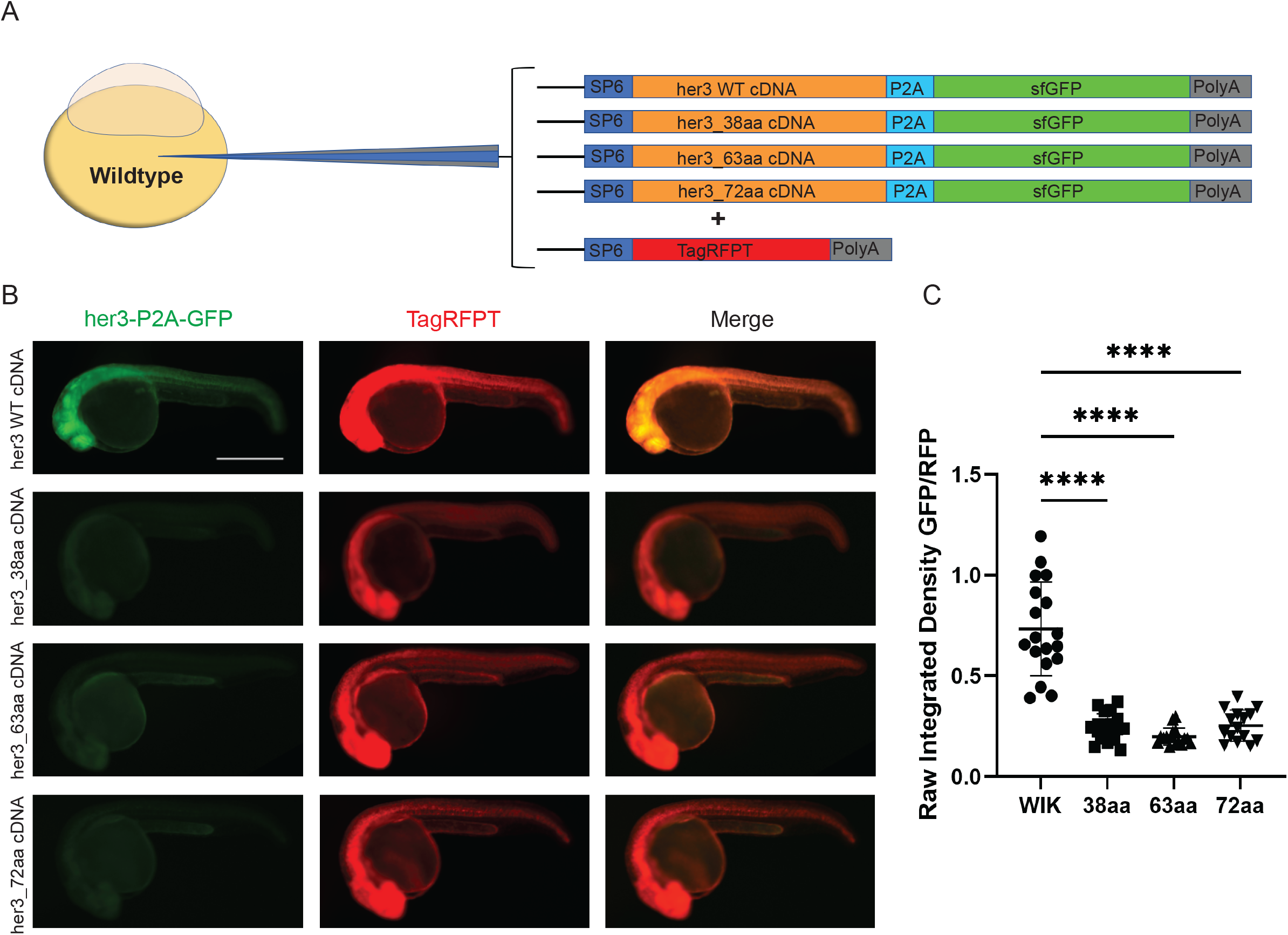
Validating *her3* frameshift mutations. (A) Schematic of strategy for *her3* mRNA expression assay. Zebrafish embryos were co-injected with a mixture of two different mRNAs: one of four different her3-P2A-sfGFP generated from either wildtype *her3* or each of the three mutant *her3* zebrafish; and a control TagRFPT mRNA. mRNA mixtures were injected into the yolk of zebrafish embryos at the 1-4 cell stage. (B) At 24 hours post fertilization (hpf), embryos were imaged on a Leica M205FA fluorescent microscope to quantify GFP and RFP fluorescence. GFP fluorescence indicates either the *her3* wildtype or *her3* mutant P2A-sfGFP construct, whereas TagRFPT is an injection control. Scale bar is 500μm. (C) Plotted is the raw integrated density ratio of GFP to RFP fluorescence. Each mark represents an individual fish. Wildtype (WIK): n=18. 38aa: n=19. 63aa: n=16. 72aa: n=15. The comparisons include wildtype WIK to *her3* mutants with the following P values: WIK-38aa: p=1.2×10^−7^. WIK-63aa: p=5.24×10^−8^. WIK-72aa: p=1.6×10^−7^.

### *her3* knockout fish are viable as adults

We then generated F2 adult homozygous fish from each of the three *her3* knockout lines. The progeny of heterozygote in-crosses had normal survival as embryos, juveniles, and adults (Figure S3). After tail clipping and genotyping sexually mature fish, and performing a HRMA for zygosity, we did not observe a significant deviation from the expected Mendelian ratios for all three potential mutants as determined by chi-square test (Figure S3). These data indicate that *her3* knockout is not lethal, which is similar to the *Hes3* mouse knockout that has been described (Hirata et al, 2001). The gross morphology of the adult fish is relatively normal, with 10% of adult fish having abnormal eye development (Figure S4). Further, homozygous *her3* mutant fish are viable and are capable of producing offspring that develop into adulthood.

### *her3* knockout embryos exhibit de-regulation of the *her3* locus and reduced overall size

A previous study of *her3* activity in zebrafish showed that *her3* directly regulates its own expression by binding to the DNA and acting as a transcriptional repressor, such that knocking down *her3* via morpholino (MO) results in an increase in *her3* mRNA expression (Hans et al, 2004). Similarly, we found that at 24hpf, *her3* knockout also results in an increase in *her3* mRNA expression in all three *her3* mutant alleles. This expression phenocopies a *her3* morpholino knock-down (Figures 3A and 3B), suggesting that the *her3* knockout is not impacted by cryptic activation that is interfacing with the *her3* locus. From this point, we decided to move forward with only the her3_38aa line because all three mutant lines were functionally similar based on the *her3* qPCR assay (Figure 3A) and because the mutation in her3_38aa produces the shortest protein product. By 24hpf, *her3* expression is spatially restricted to portions of the brain and tail as determined by fluorescent in-situ hybridization using a custom *her3* RNAscope probe, and is consistent with previous reports (Thisse and Thisse, 2005) (Figures 3C-E). In the tail, we find an expansion of *her3* mRNA in knockout fish as compared to wildtype controls (Figure 3D-E), suggesting inappropriate spatial distribution. Interestingly, in the her3_38aa mutant, we observe that the *her3* knockout embryos appear smaller than stage-matched wildtype embryos. To quantify this effect, we performed a DAPI stain of 24hpf embryos and measured the number of DAPI-positive pixels, which revealed that the *her3* knockout embryos had significantly less fluorescence (Figures 4A-C). This analysis was complemented by quantifying the overall area of the *her3* knockout embryos which was significantly reduced, indicating an impairment in developmental processes (Figure 4D).

**Figure 3:**
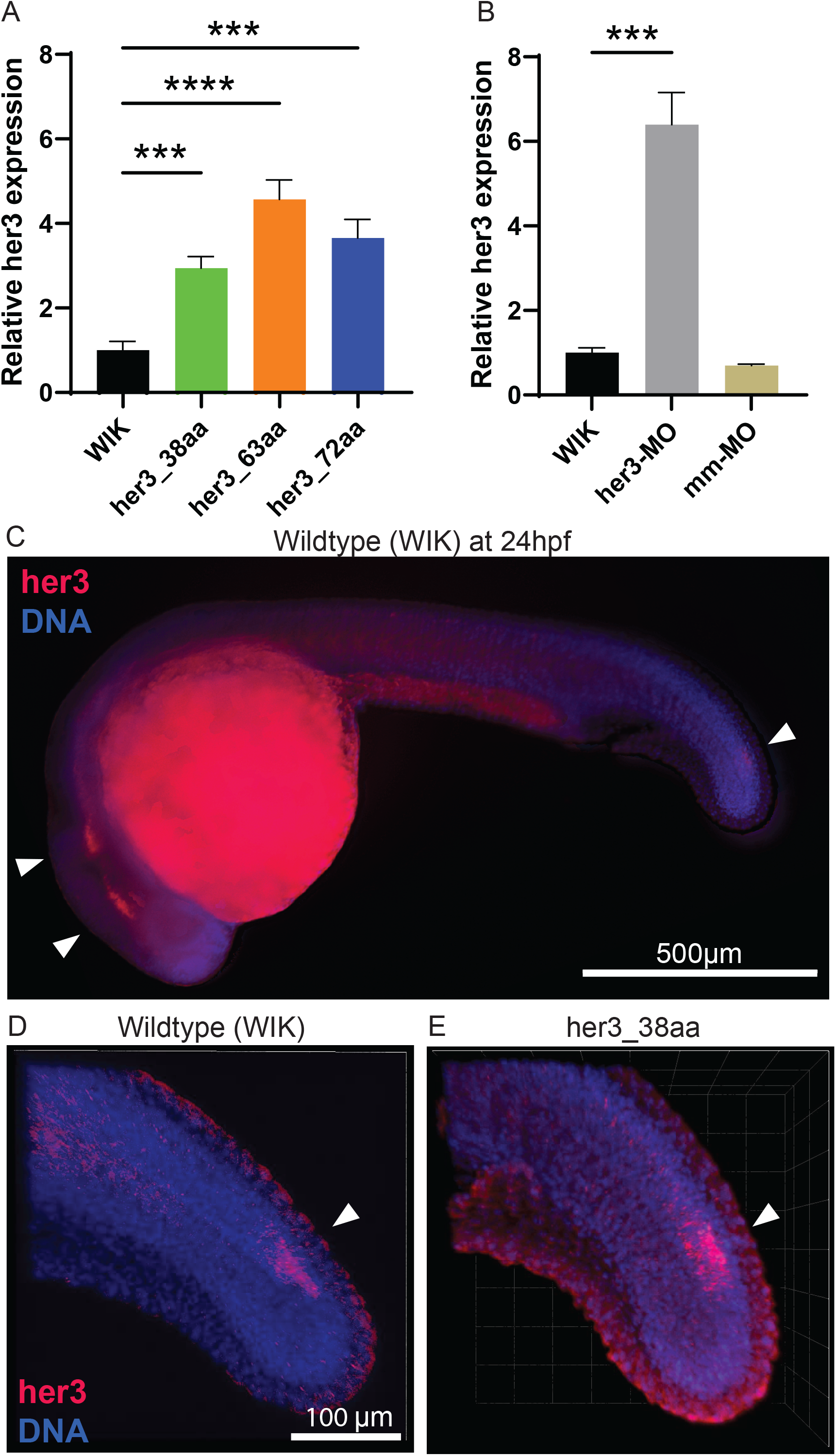
*her3* knockout recapitulates *her3-*morpholino phenotype. (A) qPCR for *her3* of each *her3* knockout line compared to the wildtype, WIK. qPCR was done with 3 biological replicates per line, with each replicate containing a pool of 10, 24 hours post-fertilization (hpf) embryos. Each biological replicate was split into three technical replicates. A Brown-Forsythe and Welch ANOVA was performed with Dunnett T3 multiple comparison test. WIK-38aa: p=1.51×10^−4^. WIK-63aa: p=6.34×10^−5^. WIK-72aa: p=6.03×10^−4^. (B) qPCR for *her3* with WIK, *her3-*MO, and mismatch (mm)-MO. 5-10 nL of morpholinos at a concentration of 500μM were injected into the yolk of WIK embryos at the 1-4 cell stage. qPCR was done with 3 biological replicates per condition, with each replicate containing a pool of 10, 24hpf embryos. Each biological replicate was split into three technical replicates. A Brown-Forsythe and Welch ANOVA was done with Dunnett T3 multiple comparison test. WIK-*her3*-MO: p=2.1×10^−4^. WIK-mm-MO: p=6.5×10^−2^, not significant. (C) Whole 24hpf zebrafish embryo. RNAscope targeting *her3* RNA transcripts was done and counterstained with DAPI. Image was taken on a Leica M205FA fluorescent stereoscope. *her3* is shown in red and DAPI is shown in blue. Scale bar is 500μm. (D-E) RNAscope targeting *her3* RNA transcripts was done on 24hpf WIK (D) and 24hpf her3_38aa (E) embryos and counterstained with DAPI. Images were taken on a Zeiss LSM800 confocal microscope with a 20x objective. Images shown are z-stack projections: (D) WIK is a projection of 53, 2μm slices; (E) her3_38aa is a projection of 48, 2μm slices. *her3* is shown in red and DAPI is shown in blue.

**Figure 4:**
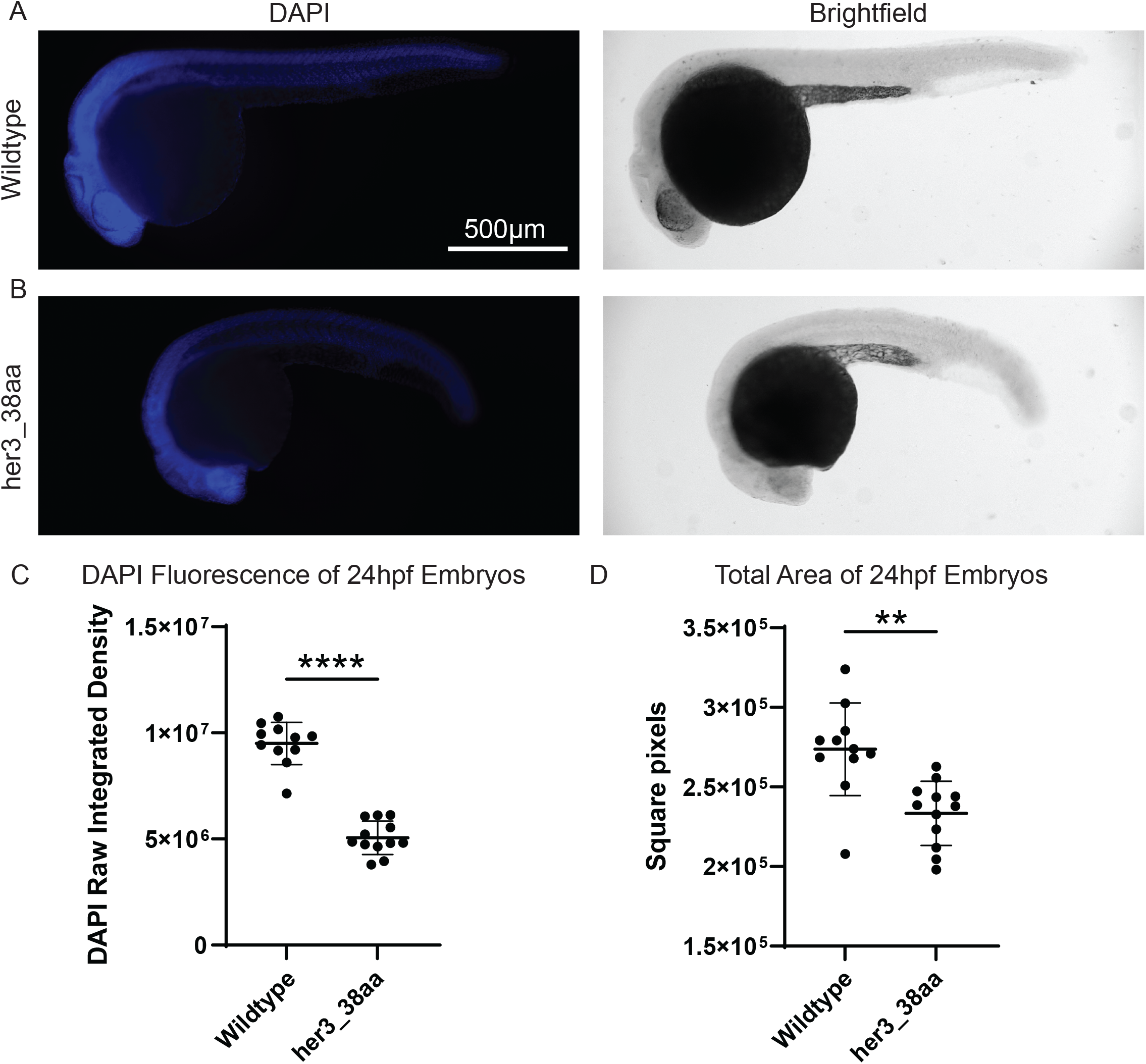
*her3* knockout line, her3_38aa, is significantly smaller than wildtype WIK at 24hpf. (A-B) DAPI and brightfield images of 24hpf wildtype WIK (A) and 24hpf her3_38aa (B) embryos. Images were taken on a Leica M205FA fluorescent stereoscope. DAPI is shown in blue. Scale bar is 500μm. (C) The raw integrated density of DAPI fluorescence was measured and graphed here. A Welch’s two-tailed T-Test was done. WIK: n=11. her3_38aa: n=12. p=3.3×10^−10^. (D) The area of the fish from the brightfield images was also measured and graphed here. A Welch’s two-tailed T-Test was done. p=1.3×10^−3^.

### *her3* knockout alters the zebrafish embryo transcriptome

To more broadly investigate the impact of *her3* loss on zebrafish development, we performed RNA-seq on her3_38aa mutants and wildtype fish at 24hpf and 72hpf. We chose these timepoints because previously we found that *her3* was up-regulated in the context of *PAX3-FOXO1* fusion-oncogene expression in 24hpf zebrafish embryos. Further, concurrent overexpression of *HES3* and *PAX3-FOXO1* increased the number of *PAX3-FOXO1* positive cells persisting between 24 and 72hpf in zebrafish embryos (Kendall et al., 2018). We also injected wildtype zebrafish with *her3*-MO and mismatch-MO and harvested injected embryos at 24hpf for comparison to the *her3* mutant (Fig. 5A). A schematic of the timepoints and groups used is in Figure 5A. Read counts can be seen in Table S1, and can be visualized along with read alignment in Figure S5A-B. For all samples, a principal component analysis (PCA) was done in which the 24hpf samples clustered together separately from the 72hpf samples, indicating that the biggest difference for demarcating groups is reflective of developmental stage. Further, the morpholino samples clustered together but separately from other 24hpf samples, suggesting there are inherent differences in uninjected and injected zebrafish. At this scale, there are fewer transcriptional changes between the *her3* mutant and the wildtype fish compared to differences between 24hpf and 72hpf (Figure 5B).

**Figure 5:**
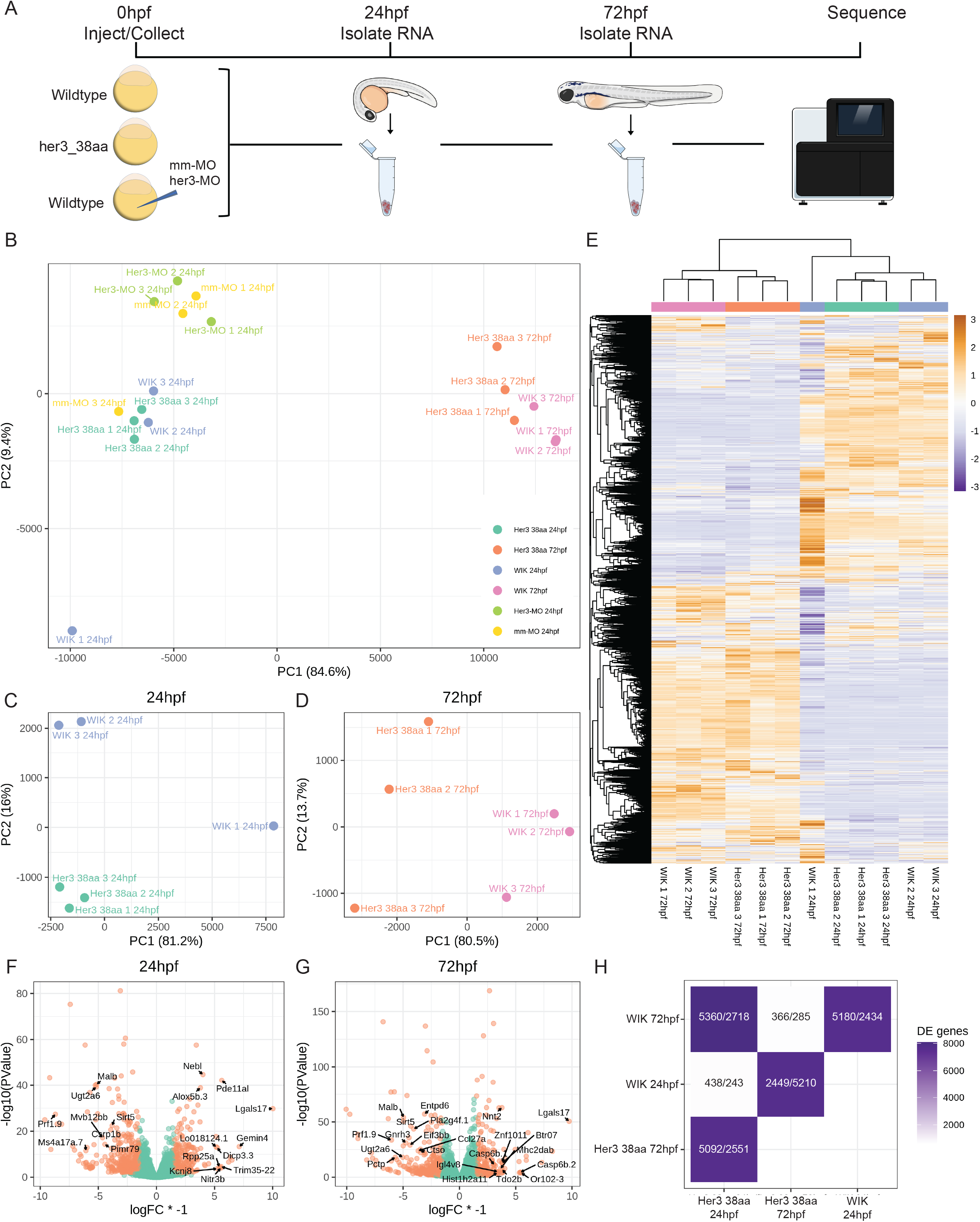
Transcriptional impact of *her3* mutation status. (A) Schematic of RNA collection. Embryos from wildtype WIK incross and her3_38aa incross were collected, with a subset of WIK embryos injected with either *her3*-MO or mismatch-MO. At 24hpf, RNA was isolated from 3 groups of 25 embryos each per condition, and again at 72hpf. RNA was then sent for sequencing. (B) PCA of all samples. (C-D) PCA of her3_38aa and WIK 24hpf samples (C) and 72hpf samples (D). (E) Heatmap clustering of all differentially expressed genes from her3_38aa and WIK samples. (F-G) Volcano plots of differentially expressed genes in her3_38aa compared to WIK 24hpf samples (F) and 72hpf samples (G). Differential expression was considered significant with an FDR ≤ 0.1 and absolute value of logFC ≥ 1.5. Top annotated up-and downregulated genes are labeled. (H) Tile chart showing pairwise comparisons of differentially expressed genes. Numbers are displayed as downregulated/upregulated of the groups along the x-axis.

However, a PCA analysis of each timepoint independently showed that her3_38aa clusters separately from WIK (Figures 5C-D). All differentially expressed genes (logFC1.5) were plotted using hierarchical clustering, indicating that WIK and her3_38aa zebrafish largely have the same expression patterns within the same timepoint (Figure 5E). We then focused on individual differentially expressed genes at 24 or 72hpf in *her3* mutants as compared to WIK visualized with volcano plots (Figures 5F-G). This highlights that the magnitude of change was amplified by 72hpf, and of interest were modulated genes such as *pctp*, which is involved in digestive tract and liver development (Zhai et al., 2017) and is significantly downregulated at 72hpf, and *csrp1b*, a cysteine-rich LIM-domain protein involved in myogenesis and fibroblast development (Iuchi and Kuldell, 2005, Bach, 2000) which is significantly downregulated at 24hpf. Figure 5H is a summary of the pairwise comparisons of up-and downregulated genes for each group, of which we focused on the *her3*-WIK 24hpf and 72hpf comparisons in Figure 6.

**Figure 6:**
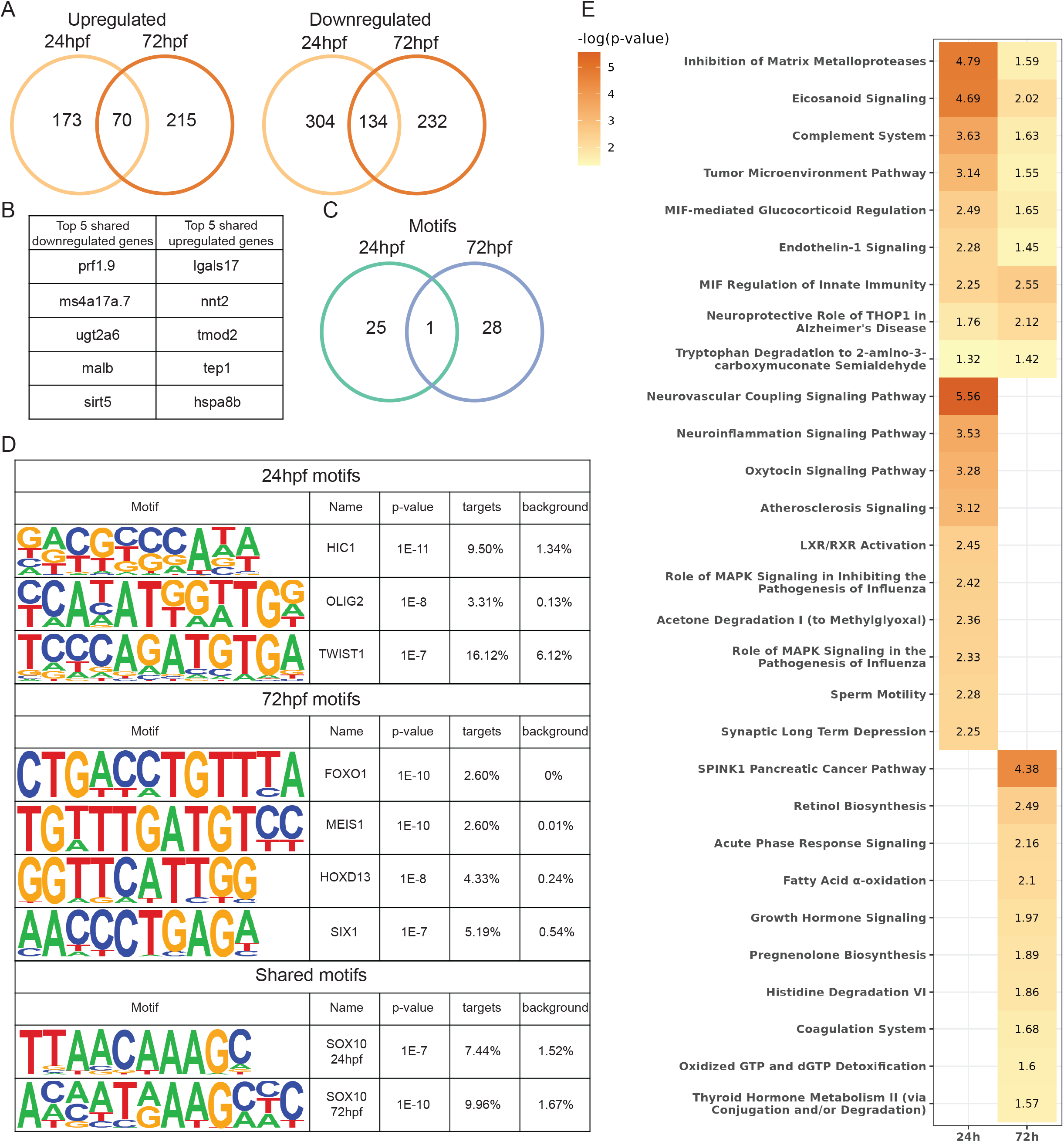
Development and cancer genes and pathways are impacted by *her3* knockout. (A) Venn diagram showing overlap of differentially expressed genes from *her3* knockout between 24hpf and 72hpf embryos. (B) List of the top 5 up-and downregulated genes shared between 24hpf and 72hpf in the *her3* knockout. (C) Venn diagram of transcription factors with DNA binding motifs overlapping between 24hpf and 72hpf *her3* knockout embryos. Targets are the percentage of differentially expressed genes that contain the motif. Background is the percentage of non-differentially expressed genes that contain the motif. (D) Examples of motifs from Venn diagram in (C). (E) p-value chart showing IPA pathways that are significantly enriched with identified differentially expressed genes.

We then assessed differentially expressed genes that were shared between the 24hpf and 72hpf timepoints for *her3* mutants as compared to wildtype embryos. These genes could suggest embryonic programs that are most impacted by the *her3* mutation or that inappropriately persist. A comparative analysis between the 24hpf and 72hpf timepoints was performed, first looking at which differentially expressed genes are shared between the two timepoints. This analysis indicates a total of 70 genes upregulated at both 24hpf and 72hpf in her3_38aa, and 134 genes that are shared and downregulated (Figures 6A-B). Shown are the annotated differentially expressed genes that are shared and unique for each of these timepoints. One gene of interest was *grinab*, downregulated at both 24hpf and 72hpf, which helps regulate apoptosis of the nervous system during development, further highlighting the mis-regulation of developmental processes in the *her3* knockout (Rojas-Rivera et al., 2012). A full list of the shared up-and downregulated genes is in Table S2. Further, a HOMER motif analysis was conducted on promoter regions of differentially expressed genes to identify transcriptional regulators outside of *her3* that are being impacted in the mutants. A single known motif was significantly enriched, IRF1, from the 72hpf samples with an adjusted p-value of 0.0304. However, 26 de novo motifs were identified in promoters of differentially expressed genes as significantly enriched in the 24hpf samples and 29 in the 72hpf samples. At 24hpf, three different *de novo* motifs for forkhead family members were identified (FOXP3, FOXQ1, and FOXH1) while FOXO1 was enriched in the 72hpf samples. A single transcription factor with DNA binding motifs was significantly enriched between 24hpf and 72hpf: sox10 (Figure 6C-D). This is perhaps not too surprising as *SOX10* is known for its role in neural crest development, but intriguingly, it is also highly expressed with functional consequences in multiple cancers such as glioma, glioblastoma, and melanoma (Pingault et al., 2022). A full list of enriched motifs is presented in Table S3.

Finally, an Ingenuity Pathway Analysis (IPA) was performed on differentially expressed genes between the *her3* mutant and wildtype embryos at both 24 hours and 72 hours post fertilization (Figure 6E). This analysis identified genes enriched within shared pathways at both timepoints, and also uniquely at the 24hpf or 72hpf timepoint. Notably, the 24hpf timepoint had enrichment for pathways involved in neuroinflammation and neural development, suggestive of *her3*’s normal role during this time period, whereas the 72hpf timepoint was enriched for SPINK1 mediated carcinogenesis. The most enriched and shared pathway between both 24hpf and 72hpf timepoints was an inhibition of matrix metallopeptidases, which is consistent with our previous data indicating that *HES3* overexpression in Rh30 *PAX3-FOXO1* rhabdomyosarcoma patient cancer cells increases *MMP3/9* expression (Kendall et al., 2018). These data correlated between the fish and human, indicating that the modulation of this pathway could have a role in ARMS pathogenesis.

## Discussion

Here, we created a zebrafish *her3* knockout line to investigate *her3’s* role in development and how these processes are potentially being co-opted in alveolar rhabdomyosarcoma. We generated a series of zebrafish *her3* mutant alleles using CRISPR/Cas9-knockout and found that these mutations were not embryonic lethal and displayed the expected Mendelian ratios. We showed that the *her3* knockout zebrafish embryos are significantly smaller than their wildtype counterparts, and that as a transcriptional repressor, removing *her3* negatively impacts appropriately controlled development. Interestingly, our RNA-seq data showed that the developmental stage had more impact on the gene expression profile than the *her3* status. However, certain genes and pathways were identified as *her3*-regulated across multiple timepoints such as an enrichment for sox10 binding motifs and regulation of genes involved in matrix metalloproteinase pathways. Taken together, these data further implicate *her3*/*HES3* in ARMS pathogenesis, with a potential role in suppressing differentiation in these immature muscle-like tumor cells.

Previously, zebrafish *her3* has been investigated in early developmental stages to understand its role in transcriptional repression. One study found that *her3* is a transcriptional repressor of *neurog1* in early development (through the 16-somite stage) and that *her3* itself is repressed by Notch signaling (Hans et al., 2004). A more recent study identified *her3* as a target of *pou5f1*, an ortholog of *OCT4*, and refined *her3’s* maximum expression to between 7-8hpf (Onichtchouk et al., 2010). We saw no significant change in *neurog1* mRNA levels at 24hpf in the *her3* knockout fish; however, this could be due to our evaluation of a later timepoint, alternative regulatory mechanisms, or potential compensation for *her3* knockout. We chose this timepoint because in our previous work, injection of the *PAX3-FOXO1* fusion-oncogene induced *her3* overexpression at 24hpf. Further, overexpression of the human *HES3* and *PAX3-FOXO1* resulted in more *PAX3-FOXO1* positive cells that inappropriately persisted by 72hpf (Kendall et al., 2018). In this study, our data provides valuable insight to the longer-term effects of the loss of a developmental transcription factor that normally possesses a finite expression window. While the number of differentially expressed genes may be few at these later timepoints (Figure 5H), the effects appear to be far reaching. In the her3_38aa mutants, reduced gene expression of *pctp, csrp1b*, and *grinab*, all genes involved in organ development, could very well explain the reduced size seen in mutant embryos and eye defects in a subset of adults. Another possibility is that *her3* has an additional function other than transcriptional repression. Recently, the bHLH transcription factor MyoD was found to be a genome organizer, helping to define the 3D structure of chromatin during myogenesis with little impact on gene expression levels (Wang et al., 2022). It is possible that the bHLH *her3/HES3* is acting in a similar manner during development, organizing chromatin in a specific 3D structure throughout the whole embryo early, and in neural tissue later in development (Thisse and Thisse, 2005). This is a direction that we plan on investigating further.

Mouse *Hes3*, the ortholog to zebrafish *her3*, is involved in the regulation of neural development and poising neural stem cells to differentiate or divide (Poser et al., 2013). Additionally, higher *Hes3* expression has been linked to increased cellular proliferation (Masjkur et al., 2014, Park et al., 2013, Poser et al., 2014). It is therefore not so surprising that human *HES3* is overexpressed in multiple cancer types with functional consequences. *HES3* was found to enhance the malignant phenotype in non-small cell lung cancer (NSCLC) and facilitated poor differentiation, increased metastasis, and worse patient prognosis. Further, *HES3* expression in NSCLC tissue upregulated cyclin D1, cyclin D3, and MMP7 (Fang et al., 2019). Our previous work found that *HES3* is a cooperating gene in *PAX3-FOXO1* fusion-driven alveolar rhabdomyosarcoma. Overexpression of *HES3* predicted reduced overall survival in patients; however, the mechanism for *HES3* and *PAX3-FOXO1* cooperation in ARMS is still unclear (Kendall et al., 2018). The similarities in *HES3* expression in NSCLC and alveolar rhabdomyosarcoma are suggestive. While cyclins are not differentially regulated in our zebrafish *her3* knockout, *mmp9* is significantly downregulated, and the inhibition of the matrix metalloproteinase pathway is significantly associated with her3 mutant transcriptional signatures (Figure 6D). Additionally, our previous work showed that HES3 overexpression in Rh30 PAX3-FOXO1 rhabdomyosarcoma patient cells upregulated *MMP3/9* expression (Kendall et al., 2018). This regulation of the matrix metalloproteinase pathway is indicative of a possible mechanism of cooperation in more aggressive cancer, as MMP3/9 facilitate metastasis (Nagase et al., 2006). We also found SOX10 binding motifs to be significantly enriched for among the differentially expressed genes in the *her3* knockout, which is suggestive of how *her3/HES3* may be functioning in these cancers (Figure 6D). A recent study in *Xenopus* found that overexpression of *hes3* resulted in an increase in *sox10* expression, which caused the neural crest progenitors to maintain an undifferentiated state (Hong and Saint-Jeannet, 2018). In addition to SOX10 motifs, several other motifs were identified that further links *her3* to cancer (Figure 6D). HIC1 and TWIST1 have been implicated in breast cancer (Wang et al., 2018, Martin et al., 2005). OLIG2 is highly expressed in gliomas (Ligon et al., 2004). SIX1 has been shown to regulate cancer stem cell characteristics and numbers (Kingsbury et al., 2019). The enrichment of DNA binding motifs for these cancer-related genes is indicative of the strong link *her3* has to cancer contexts.

Overall, our new *her3* zebrafish knockout is a powerful model with dual utility to study development, cancer, and the consequences of their interface. Given that the mechanism of action of *HES3* in ARMS is still unclear, we plan to use this model to study how the cancer-driver fusion protein *PAX3-FOXO1* interacts with and without *her3/HES3*. By complementing these studies with the analysis of patient tumors and cell culture models, we anticipate that this model will help further elucidate how *her3/HES3* is acting in both development and cancer, which are intrinsically linked.

## Methods

### Zebrafish Humane Use and Husbandry

Zebrafish are housed in an AAALAC-accredited, USDA-registered, OLAW-assured facility in compliance with the Guide for the Care and Use of Laboratory Animals. All research procedures are approved by the IACUC at The Abigail Wexner Research Institute at Nationwide Children’s Hospital according to IACUC protocol AR19-00172. Zebrafish are free of Pseudoloma neurophilia, Pleistophora hyphessobryconis, Pseudocapillaria tomentosa, Mycobacterium spp., Edwardsiella ictalurid, Ichthyophthirius multifilis, Flavobacterium columnare, and zebrafish picornavirus (ZfPV1) as determined by quarterly sentinel monitoring program. The fish were housed at a density of 8-12 fish per liter in mixed-sex groups in 0.8 L, 1.8 L, 2.8 L, or 6 L tanks on a recirculating system (Aquaneering, San Diego, CA) in 28ºC water (conductivity, 510 to 600 μS; pH, 7.3 to 7.7; hardness, 80 ppm; alkalinity, 80 ppm; dissolved oxygen, greater than 6 mg/L; ammonia, 0 ppm; nitrate, 0 to 0.5 ppm; and nitrite, 0 ppm) in a room with a 14:10-h light:dark cycle. System water was carbon-filtered municipal tap water, filtered through a 20-μm pleated particulate filter, and exposed to 40W UV light. The fish were fed twice daily with both a commercial pelleted diet and a live brine shrimp cultured in-house. WIK were used as the wildtype line and were obtained from the Zebrafish International Resource Center (ZIRC; https://zebrafish.org). *her3* mutant zebrafish will be made available upon request and will be deposited at ZIRC.

### Cloning

pCS2+MCS-P2A-sfGFP was a gift from Jason Berman (Addgene plasmid # 74668; http://n2t.net/addgene:74668; RRID:Addgene_74668) (Prykhozhij et al., 2017). pCS2-TagRFPT.zf1 was a gift from Harold Burgess (Addgene plasmid # 61390; http://n2t.net/addgene:61390; RRID:Addgene_61390) (Horstick et al., 2015). RNA was collected from zebrafish embryos from either wildtype WIK or homozygous *her3*-knockout using QIAGEN RNeasy Mini kit (74104, QIAGEN). Then, cDNA was synthesized using the RT2 HT First Strand Synthesis kit (330411, QIAGEN). PCR was done using the cDNA and primers against the *her3* coding sequence (Table S4), adding a PacI site on the 5’ end and an AscI site on the 3’ end. PCR products were cleaned up using the Monarch PCR Cleanup kit (T1030L, NEB). A double digest using PacI and AscI was done on the PCR products and the pcs2+MCS-P2A-sfGFP plasmid, and then cleaned up using the Monarch PCR Cleanup kit (T1030L, NEB). Digested PCR products were then ligated to the digested pcs2+MCS-P2A-sfGFP plasmid using T4 DNA ligase (M0202S, NEB) in a 3:1 insert:backbone ratio. Ligated plasmids were transformed into DH5α cells (FEREC0111, Fisher Scientific). Colonies were picked and sequenced via Sanger sequencing by Eurofins Genomics to verify insert. Final plasmids were named pcs2+WTher3-P2A-sfGFP, pcs2+her3_38-P2A-sfGFP, pcs2+her3_63-P2A-sfGFP, and pcs2+her3_72-P2A-sfGFP.

### Zebrafish Embryo Injections

Zebrafish were injected at either the 1-cell stage in the cell body for CRISPR/Cas9 injections, or the 1-4 cell stage in the yolk for RNA and morpholino injections. For CRISPR/Cas9 injections, embryos were injected with 36ng/μl crRNA and 67ng/μl of tracrRNA for one of three gRNAs against the *her3* locus (Table S4), 100 ng/μl Cas9 protein (IDT), 0.1% phenol red, and 0.3X Danieau’s buffer. For RNA injections, embryos were injected with: 100 ng/μl RNA of either pcs2+WTher3-P2A-sfGFP, pcs2+her3_38-P2A-sfGFP, pcs2+her3_63-P2A-sfGFP, or pcs2+her3_72-P2A-sfGFP; 50ng/μl RNA of pcs2-tagRFPT.zf1; 0.1% phenol red; and 0.3X Danieau’s buffer. For morpholino injections, embryos were injected with: 200 μM of either *her3*-MO1 or mm-MO (Table S4); 0.1% phenol red; and 0.3X Danieau’s buffer. Morpholinos were custom ordered from GeneTools.

### High Resolution Melt Analysis

CRISPR knockouts were verified using High Resolution Melt Analysis (HRMA). Either embryos or fin clips were used to generate genomic DNA for HRMA. A minimum of three wildtype (WIK) samples were included with each HRMA as the reference group. Briefly, individual 24hpf embryos or adult fin clips were incubated in 10 mM Tris-HCl pH 8.3, 50mM KCl, 0.3% Triton X-100, and 0.3% NP40 for 10 minutes at 98°C, then cooled on ice for 10 minutes. Then, 1/10^th^ volume of 10 mg/ml Proteinase K (BP1700, Fisher Scientific) was added and samples were incubated overnight at 55°C, heated to 98°C for 10 minutes to heat-inactivate proteinase K, and cooled to between 4°C and 12°C. For HRMA, 200 ng of genomic DNA, 9 μl of Precision Melt Mix (1725112, Bio-Rad), 0.5 μl of each of forward and reverse primers (Table S4), and water up to 20 μl was added to a 384-well PCR plate. The plate was then run on a Bio-Rad CFX384 Real Time PCR Detection System using the following program: 95°C for 3 minutes; (95°C for 15 seconds, 60°C for 20 seconds, 70°C for 20 seconds) x45; 65°C for 30 seconds; melt 65°C-95°C, 0.2°C/step hold 5 seconds; 95°C for 15 seconds. The real-time data file was then opened on Bio-Rad Maestro software to generate a .PCRD file. The .PCRD file was then opened on Bio-Rad Precision Melt Analysis software. Known wildtype WIK samples were used as the reference group. Samples with a Difference RFU between 0.1 and -0.1 were excluded.

### Real Time Quantitative PCR

Real time qPCR was done on both 24hpf and 72hpf zebrafish embryos. Briefly, 3 pools of 10 embryos from each of WIK, her3_38aa, her3_63aa, and her3_72aa at both 24hpf and 72hpf were collected. Additionally, 3 pools of 10 embryos each of WIK embryos injected with either *her3*-MO1 or mm-MO (Table S4) at 24hpf were collected. RNA was isolated from each of the pools of embryos using the QIAGEN RNeasy Mini kit (74104, QIAGEN) including an on-column DNAse digestion. The RT2 HT First Strand Synthesis kit (330411, QIAGEN) was used to synthesize cDNA using 160 ng of RNA per sample. cDNA was diluted to 80μl with nuclease-free water. A mastermix containing 5 μl SYBR Green 2x (1725122, Bio-Rad) and 0.5 μl of each 10 mM forward and reverse primer, per sample, was combined with 4 μl of cDNA in a 384-well PCR plate. The plate was run on a CFX384 Real Time PCR Detection System using the following program: 95°C for 2 minutes; (95°C for 15 seconds, 60°C for 1 minute, Plate Read) x40; 65°C for 30 seconds; (65°C, 0.5°C/step, Plate Read) x60. The real time data file was then opened on Bio-Rad CFX Maestro software to generate a .PCRD file. The software was then used to determine differential gene expression using the ΔΔCT method. Housekeeping genes used: *rpl13a* and *actinb1* (Table S4). Genes queried: *her3* (Table S4).

### RNAscope and Nuclear Staining

For RNAscope, 24hpf zebrafish embryos from both WIK and her3_38aa homozygous in-cross were collected. The RNAscope Assay on Whole Zebrafish Embryos as well as the RNAscope Multiplex Fluorescent v2 (323100, Advanced Cell Diagnostics) manuals were followed. The dre-her3 probe (895001-C2, Advanced Cell Diagnostics) was used with DAPI provided in the Multiplex Fluorescent kit. Opal 570 (NC1601877, Fisher Scientific) was used. Embryos were imaged on both a Zeiss LSM800 inverted confocal microscope with a 20x objective and a Leica M205FA fluorescent stereoscope with a 2x objective. For immunofluorescence, 24hpf embryos from both WIK and her3_38aa were collected and fixed in 4% paraformaldehyde (PFA) for 2 hours at room temperature. Embryos were then dehydrated in 50% methanol for 5 minutes and rinsed twice with 100% methanol. Embryos were then stored in 100% methanol at -20°C for 30 minutes to up to 6 months. Embryos were rehydrated in 50% methanol for 5 minutes, then washed with PBST (1x PBS with 0.1% Tween-20) for 3×5 minutes. Next, embryos were permeabilized for 1 minute in 10 μg/ml of proteinase K and rinsed three times in PBST. Embryos were then fixed for 20 minutes at room temperature in 4% PFA and rinsed 3×5 minutes in PBST. Embryos were next incubated in DAPI (ab228549, Abcam), diluted 1:1000 in PBS, for 30 minutes at room temperature. Embryos were rinsed 3×5 minutes in PBS. Embryos were imaged on a Leica M205FA stereoscope with a 2x objective. Image analysis and quantification of area and DAPI positive pixels were performed in ImageJ version 1.53h.

### RNA-seq

RNA-seq was performed on both 24hpf and 72hpf embryos. Three pools of 20 embryos each of WIK, her3_38aa, WIK injected with *her3*-MO1, and WIK injected with mm-MO at 24hpf, and of WIK and her3_38aa at 72hpf, were staged and then collected. RNA was immediately isolated using the QIAGEN RNeasy Mini kit with on-column DNAase digestion (74104, QIAGEN). RNA integrity and sample QC was determined by the Nationwide Children’s Hospital Institute for Genomic Medicine (IGM) using an Agilent Bioanalyzer 2100. WIK 24hpf samples were spiked with 2 μl of 1:100 diluted ERCC spike-in mix 1 (4456740, Thermofisher Scientific) while all other samples were spiked with 2 μl of 1:100 diluted ERCC spike-in mix 2 (4456740). IGM generated sequencing libraries using the NEBNext Ultra II Directional RNA Library Prep Kit for Illumina (E7760, New England Biolabs). A polyA enrichment step (E7490, New England Biolabs) was used to enrich for polyadenylated RNAs. RNA samples were run on a NovaSeq Sp with 150bp paired-end reads and a total of 43 – 51 million reads generated output per sample (Table S1, Figure S5).

### RNA-seq Analysis

All RNA-seq analyses were performed on the Nationwide Children’s Hospital high-performance cluster. The cluster is running CentOS 8.1 with a slurm job manager. All code used in analyzing the RNA-seq data is freely available online (https://github.com/MVesuviusC/kentRNAseq1). We used FASTQC (v0.11.9) to evaluate sequence data quality. We created a dual reference containing both the zebrafish genome (danRer11) as well as the ERCC spike-in reference sequences to use for alignment with HiSat2 (v 2.2.1) (Kim et al., 2019). We allowed for only a single alignment per read (-k 1), and then removed PCR duplicates and removed any improperly paired reads with SAMtools (v1.10) (Li et al., 2009). To count the number of reads per gene, we employed the featureCounts program from Subread (v2.0.2) package (Liao et al., 2014). We performed statistical analysis of differential expression using the EdgeR R package (v 3.34.0) (Robinson et al., 2010). To call differentially expressed genes we used a FDR cutoff of less than or equal to 0.1 and a fold-change cutoff of either greater than or equal to 1.5 or less than or equal to -1.5. For motif analysis, we used HOMER (v4.11.1) (Heinz et al., 2010). Plotting and data manipulation were primarily done using the tidyverse suite of R packages (Wickham et al., 2019).

### Statistics

GraphPad Prism 9 and R were used for all statistical analysis. A chi-square test was used to determine if observed zebrafish clutches were significantly different from expected Mendelian ratios (Figure S3). A one-way Brown-Forsythe and Welch ANOVA with a Dunnett T3 test was used to determine significance of *her3* expression (Figure 3A-B). A two-tailed Welch’s T-Test was done to determine significance of DAPI fluorescence and area of zebrafish embryos (Figure 4C-D). A two-way ANOVA with Fisher’s LSD test was done to determine significance of gene expression between WIK and her3_38aa zebrafish embryos (Figure 5A-B). All sample sizes are included in figure legends.

## Data and Code Availability

All data is freely available through the NCBI GEO repository (submission in progress) and all code is available online (https://github.com/MVesuviusC/kentRNAseq1).

## Acknowledgements

We thank Dr. Raphael Malbrue, Logan Fehrenbach, Adewole Adekanye, and the other Animal Resources Core Zebrafish Facility team members for exceptional care and collaboration in maintaining our zebrafish. We thank Dr. Stephen Lessnick and Jack Kucinski for helpful discussion. We thank the Institute for Genomic Medicine, the NCH Morphology Core, and the NCH High Performance Computing group for providing services to support this study.

## Author Contributions

Conceptualization: M.K., G.K.; Methodology: M.K., C.L., G.K.; Software: M.C.; Validation: M.K., M.C., G.K.; Formal analysis: M.K., D.C., M.C., G.K.; Investigation: M.K., D.C., K.S., C.L., G.K.; Resources: G.K.; Data Curation: M.C., G.K.; Writing – original draft: M.K., M.C., G.K.; Writing – review and editing: M.K., D.C., K.S., C.L., M.C., G.K.; Visualization: M.K., M.C., G.K.; Supervision: G.K.; Project administration: G.K.; Funding acquisition: G.K.

## Funding Sources

This work was supported by an Alex’s Lemonade Stand Foundation “A” Award, a V Foundation for Cancer Research V Scholar Grant, and Startup Funds from The Abigail Wexner Research Institute at Nationwide Children’s Hospital to G.C.K. D.C. is supported by a Graduate Enrichment Fellowship from The Ohio State University. The Institute for Genomic Medicine is funded by the Nationwide Foundation Pediatric Innovation Fund and the Ohio State University Comprehensive Cancer Center grant P30 CA016058.

## Competing Interests

The authors declare no competing or financial interests.

## Notes

https://github.com/MVesuviusC/kentRNAseq1

### Competing Interest Statement

The authors have declared no competing interest.

