## Supplemental Figures for "Zebrafish *her3* knockout impacts developmental and cancer-related gene signatures"

### Running Title: Zebrafish *her3* knockout

Matthew R. Kent[1], Delia Calderon[1,2], Katherine M. Silvius[1], Collette A. LaVigne[3], Matthew V. Cannon[1], Genevieve C. Kendall[1,2,4#]

1. Center for Childhood Cancer & Blood Diseases, The Abigail Wexner Research Institute, Nationwide Children's Hospital, Columbus, OH 43205, USA.
2. Molecular, Cellular, and Developmental Biology Ph.D. Program, The Ohio State University, Columbus, OH 43210, USA.
3. Department of Molecular Biology, UT Southwestern Medical Center, Dallas, TX 75390, USA.
4. Department of Pediatrics, The Ohio State University College of Medicine, Columbus, OH 43205, USA.

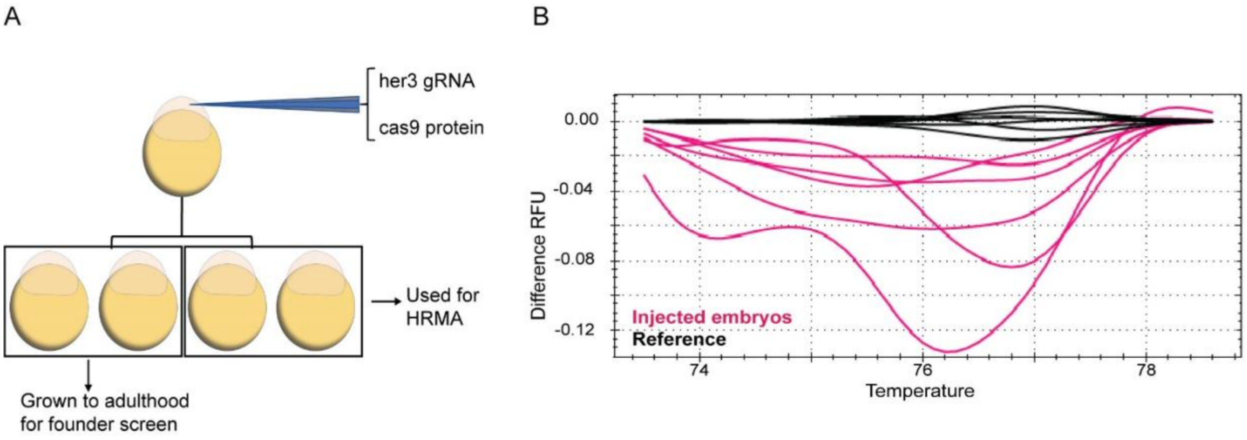


**Figure S1. Generating *her3* knockout founders via CRISPR.** (A) Schematic of injection method. Wildtype WIK embryos were injected into the cell body at the 1-2 cell stage with Cas9 protein and *her3* gRNA. A subset of embryos was collected at 24hpf to test for gRNA cutting efficiency by HRMA (B). Potential $P_0$ mutants are shown in pink.

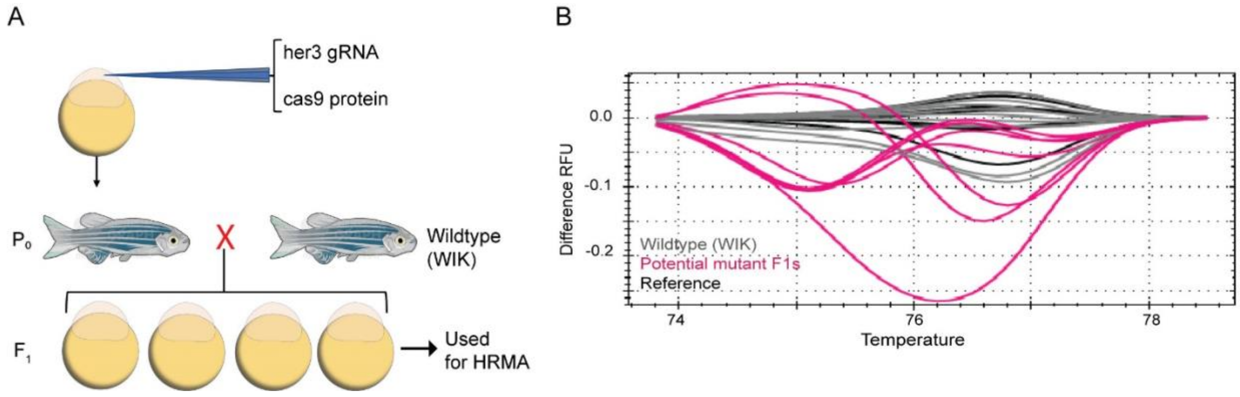


**Figure S2. Identifying *her3* knockout founders via HRMA.** (A) Schematic for founder identification strategy. Adult potential founders were outcrossed to wildtype WIK in single mating pairs. Genomic DNA was collected from single embryos and used to do HRMA (B). Potential $F_1$ mutants are shown in pink.

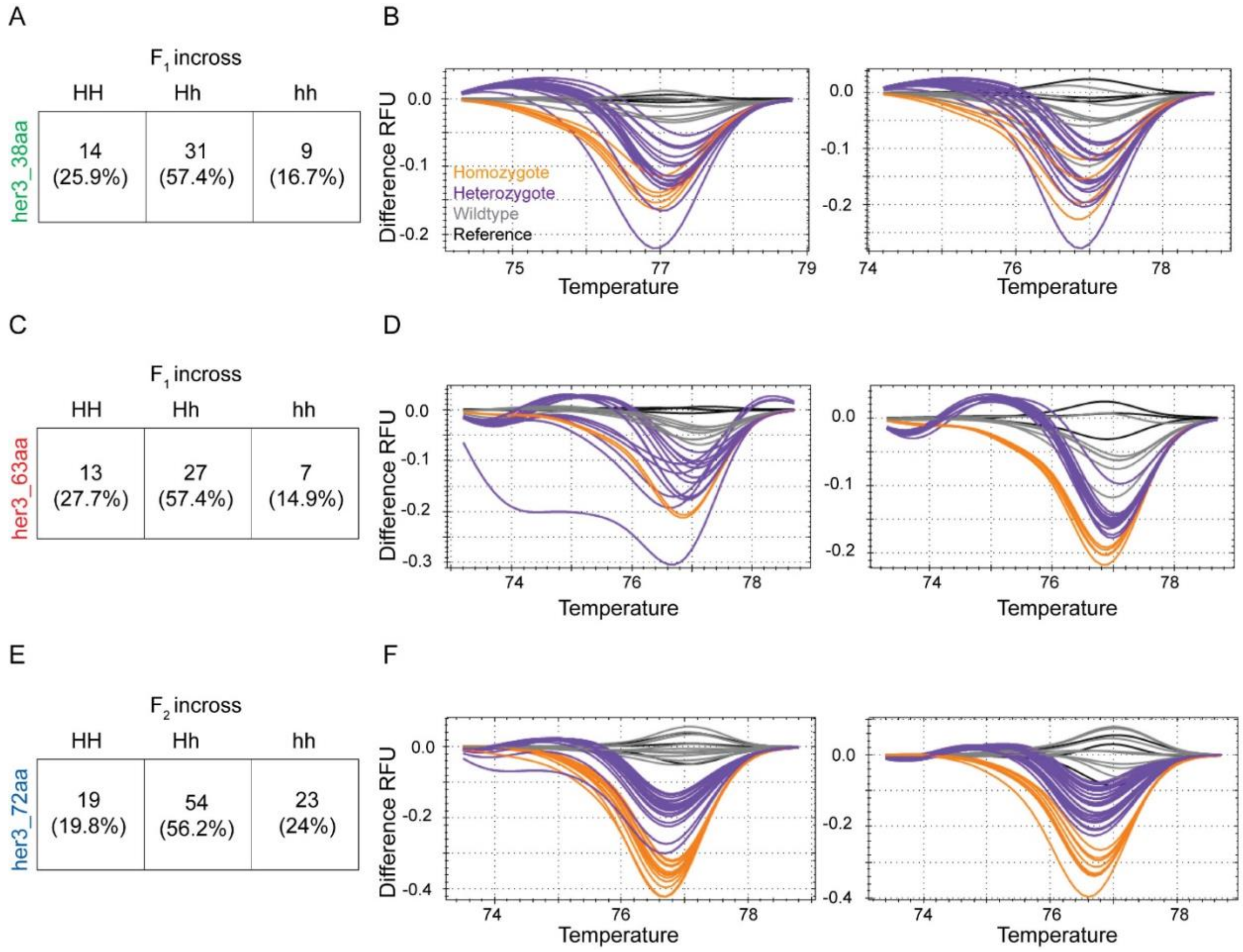


**Figure S3. *her3* knockout mutation display Mendelian ratios.** (A, C, E) Charts showing counts and percentages of each genotype from two separate heterozygous incrosses for her3_38aa (A), her3_63aa (B), and her3_72aa (C). A chi-square test was performed. (A) chi-square=2.1111, p=0.3480. (B) chi-square=2.5745, p=0.2760. (C) chi-square=1.8333, p=0.3998. HRMA was used to genotypes from each heterozygous incross (B, D, F). Fin-clips of adult progeny were taken for genomic DNA isolation. HRMA was then performed. Homozygotes are shown in orange, heterozygotes are shown in purple, wildtype are shown in gray, and reference WIK samples are shown in black.

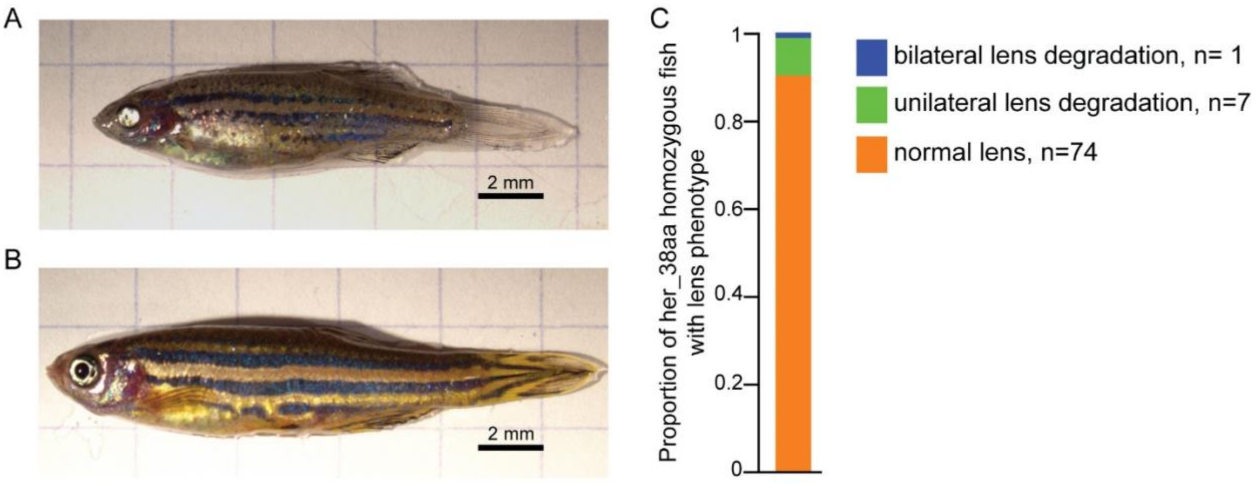


**Figure S4. A subset of *her3* knockout fish display eye defects in adulthood.** A subset of her3_38aa homozygous fish presented with eye defects after >6 months of age. Shown are representative brightfield images of a her3_38aa zebrafish that have eye defects (A) or sibling her3_38aa control with normal eyes (B). Scale bar is 2mm. (C) Quantification of the penetrance of the lens phenotype in adult fish. All fish counted were between 8 and 10 months old when lens degradation was identified. Normal lens: n=74. Unilateral lens degradation: n=7. Bilateral lens degradation: n=1.

A

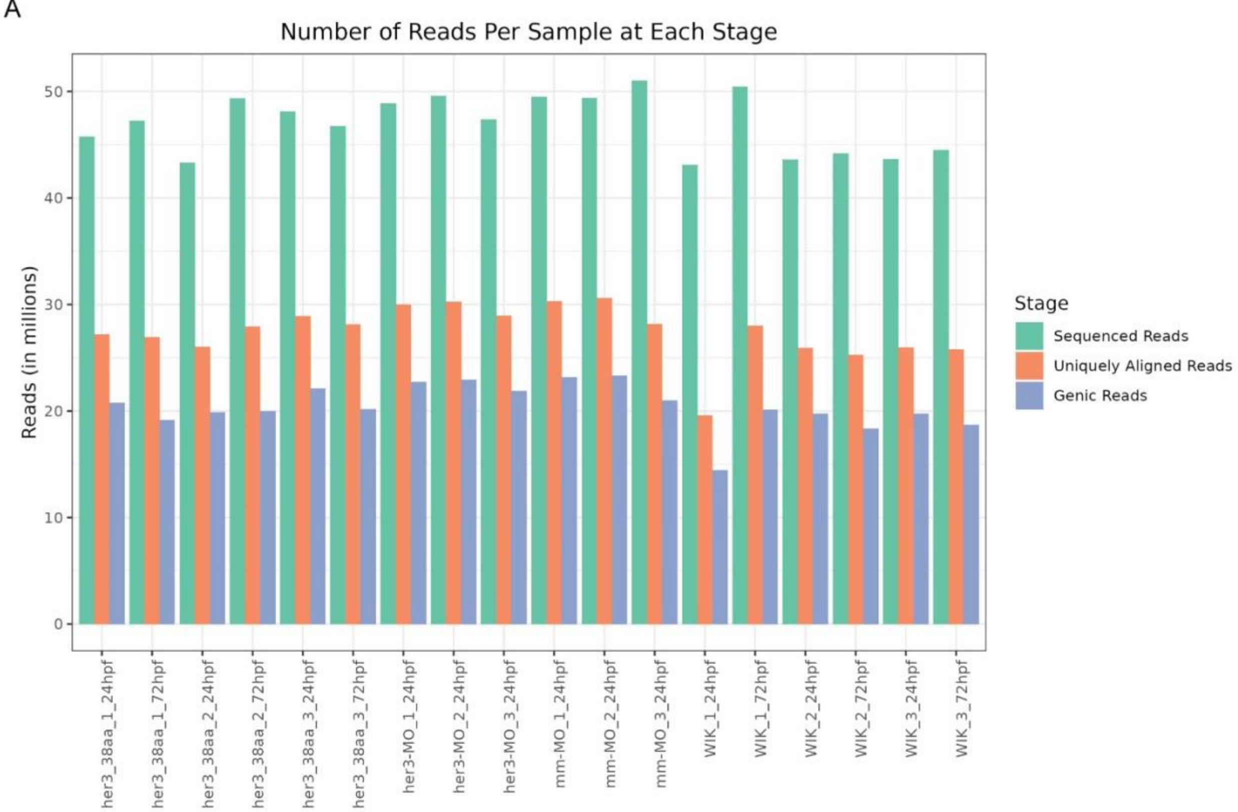
Number of Reads Per Sample at Each Stage
Reads (in millions)
Stage
Sequenced Reads
Uniquely Aligned Reads
Genic Reads
her3_38aa_1_24hpf
her3_38aa_1_72hpf
her3_38aa_2_24hpf
her3_38aa_2_72hpf
her3_38aa_3_24hpf
her3_38aa_3_72hpf
her3-MO_1_24hpf
her3-MO_2_24hpf
her3-MO_3_24hpf
mm-MO_1_24hpf
mm-MO_2_24hpf
mm-MO_3_24hpf
WIK_1_24hpf
WIK_1_72hpf
WIK_2_24hpf
WIK_2_72hpf
WIK_3_24hpf
WIK_3_72hpf

B

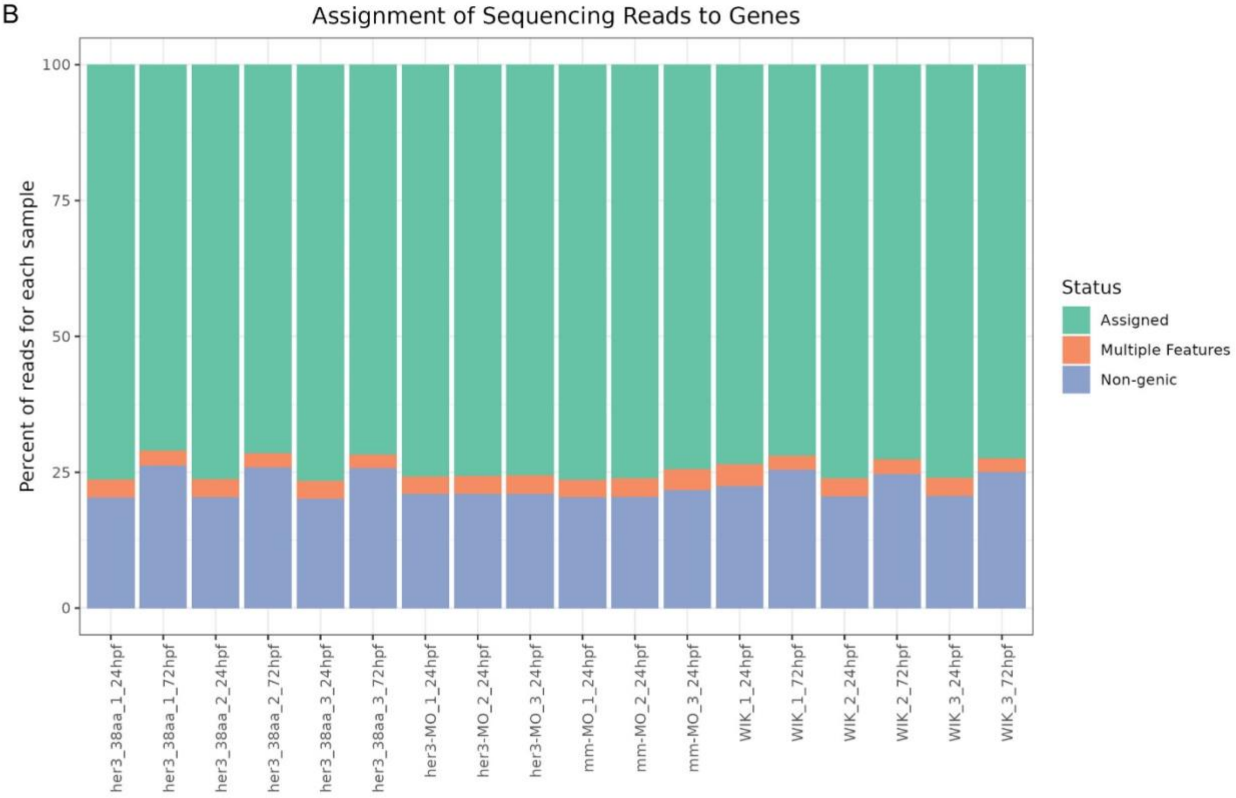
Assignment of Sequencing Reads to Genes
Percent of reads for each sample
Status
Assigned
Multiple Features
Non-genic
her3_38aa_1_24hpf
her3_38aa_1_72hpf
her3_38aa_2_24hpf
her3_38aa_2_72hpf
her3_38aa_3_24hpf
her3_38aa_3_72hpf
her3-MO_1_24hpf
her3-MO_2_24hpf
her3-MO_3_24hpf
mm-MO_1_24hpf
mm-MO_2_24hpf
mm-MO_3_24hpf
WIK_1_24hpf
WIK_1_72hpf
WIK_2_24hpf
WIK_2_72hpf
WIK_3_24hpf
WIK_3_72hpf

**Figure S5. Read alignment from RNA sequencing data.** (A) Read (in millions) per sample for sequenced reads (green), uniquely aligned reads (orange), and genic reads (blue). (B) Assignment of sequencing reads to genes. Assigned reads are shown in green, reads with multiple features are shown in orange, and non-genic genes are shown in blue.

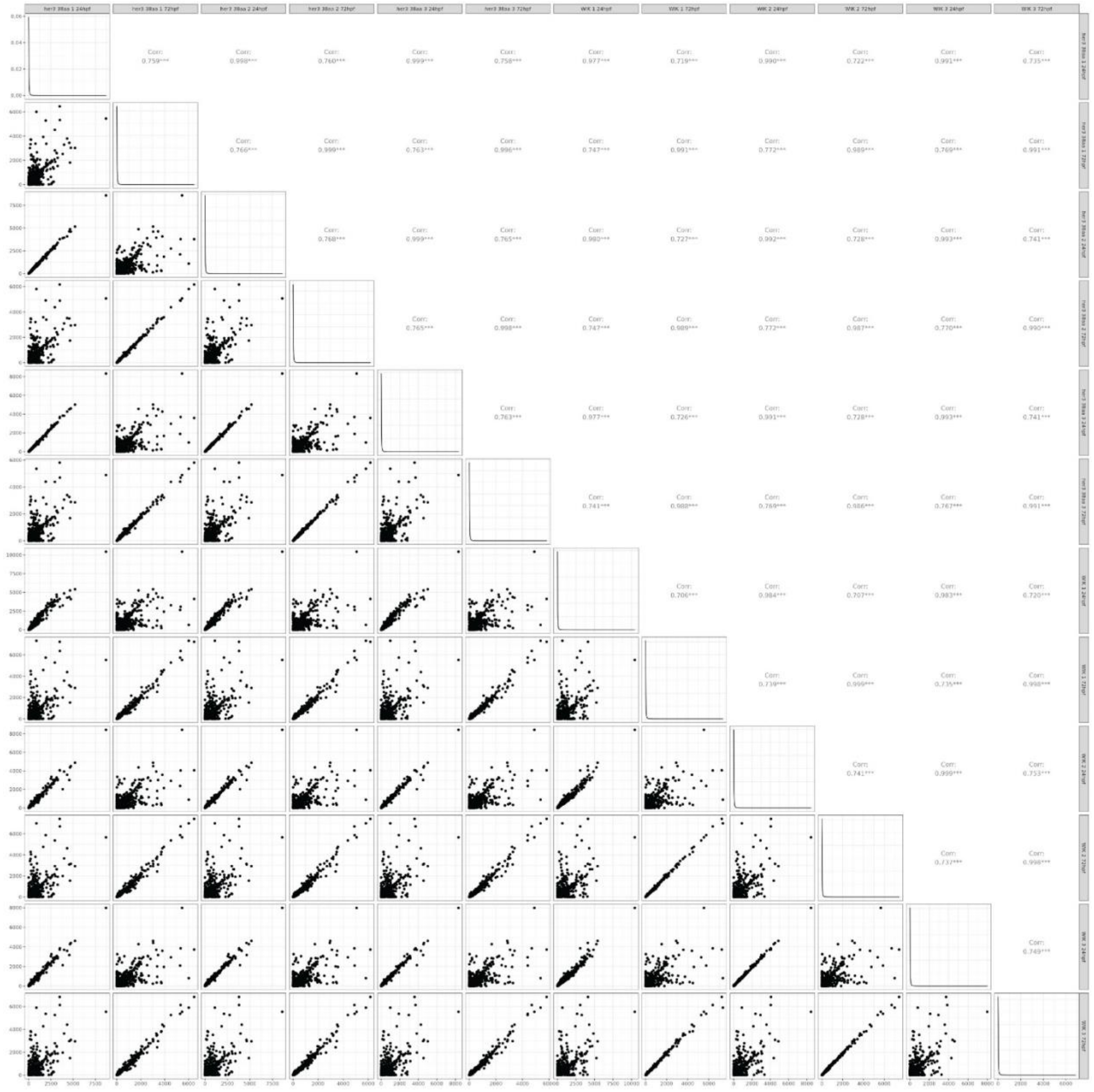

**Figure S6. her3_38aa correlates more with WIK within the same time point than with her3_38aa from a different time point.** Pairwise correlation of each RNAseq sample.

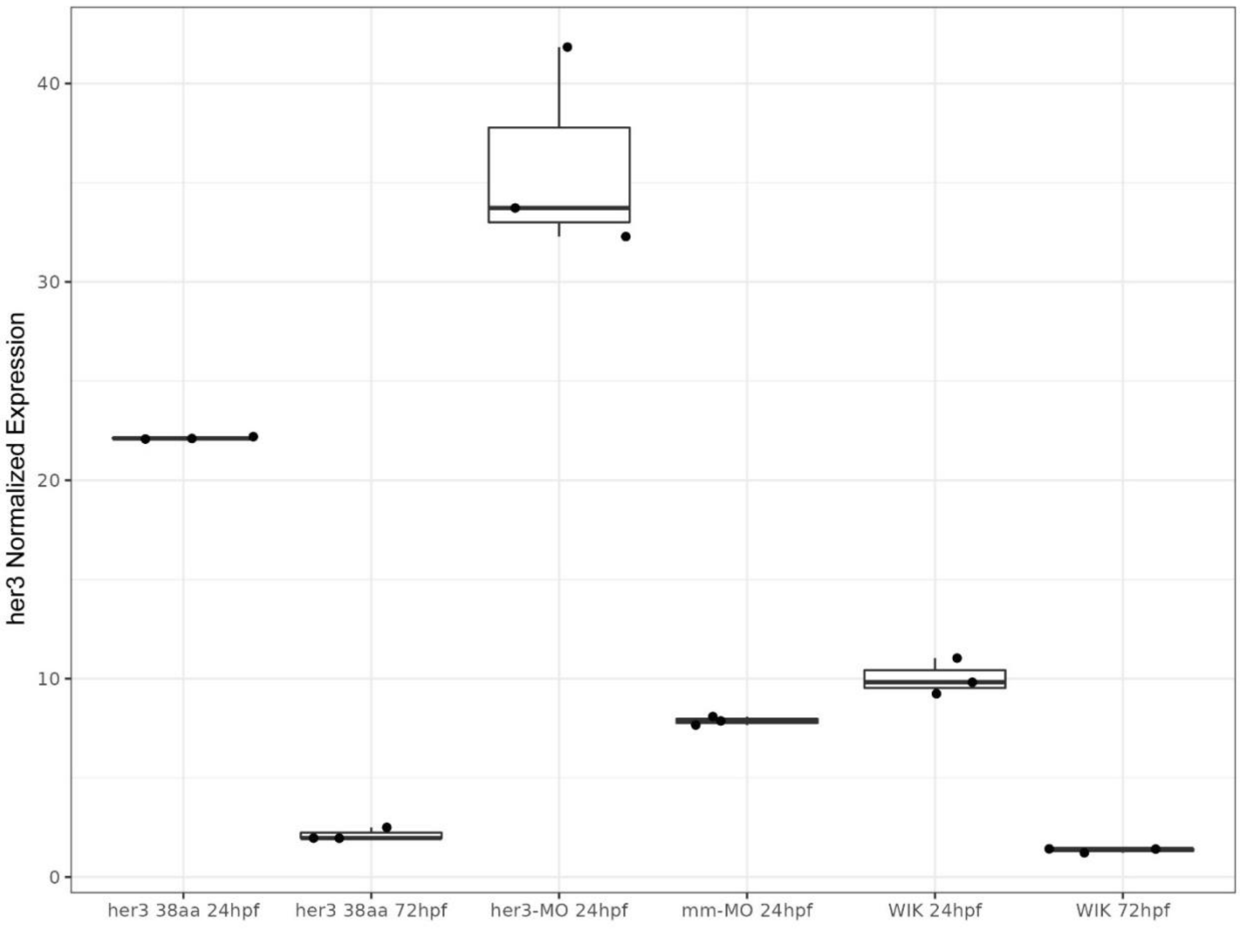


**Figure S7. Increased *her3* expression via knockout is consistent with *her3*-MO injection.** Normalized read counts of *her3* transcripts per sample group.

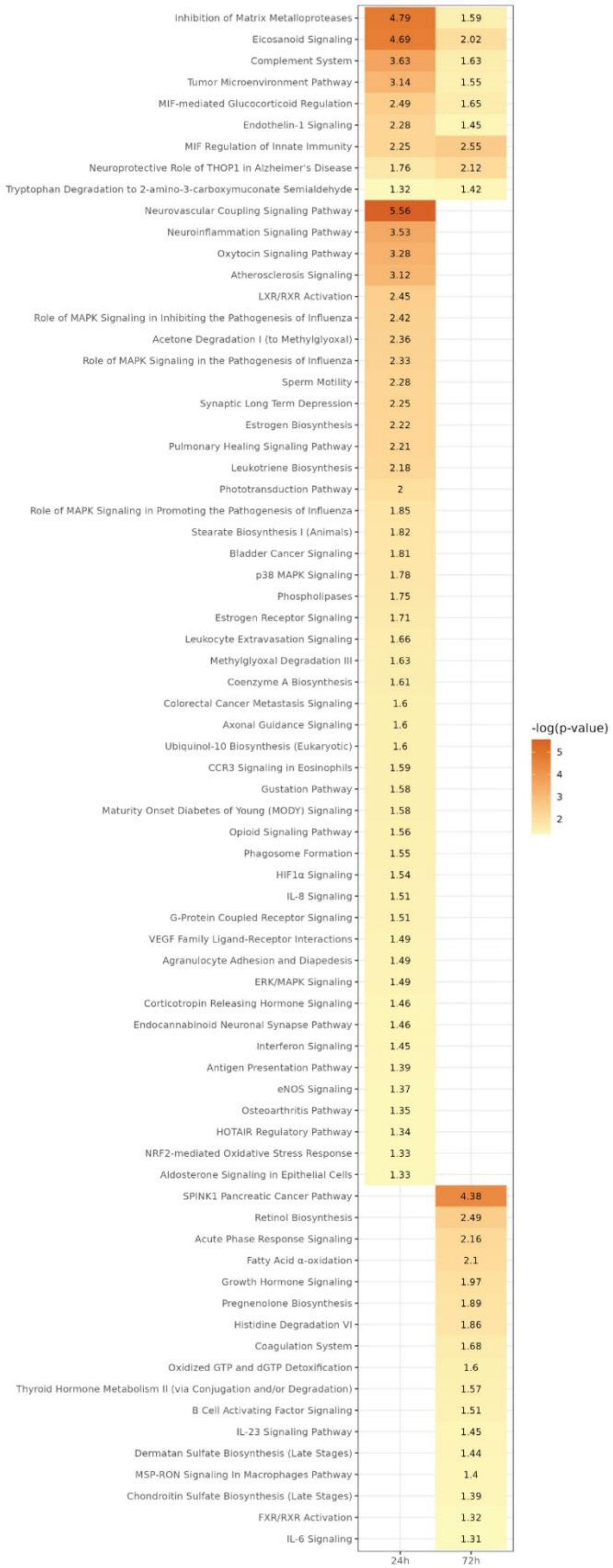
Inhibition of Matrix Metalloproteases 4.79 1.59
Eicosanoid Signaling 4.69 2.02
Complement System 3.63 1.63
Tumor Microenvironment Pathway 3.14 1.55
MIF-mediated Glucocorticoid Regulation 2.49 1.65
Endothelin-1 Signaling 2.28 1.45
MIF Regulation of Innate Immunity 2.25 2.55
Neuroprotective Role of THOP1 in Alzheimer's Disease 1.76 2.12
Tryptophan Degradation to 2-amino-3-carboxymuconate Semialdehyde 1.32 1.42
Neurovascular Coupling Signaling Pathway 5.56
Neuroinflammation Signaling Pathway 3.53
Oxytocin Signaling Pathway 3.28
Atherosclerosis Signaling 3.12
LXR/RXR Activation 2.45
Role of MAPK Signaling in Inhibiting the Pathogenesis of Influenza 2.42
Acetone Degradation I (to Methylglyoxal) 2.36
Role of MAPK Signaling in the Pathogenesis of Influenza 2.33
Sperm Motility 2.28
Synaptic Long Term Depression 2.25
Estrogen Biosynthesis 2.22
Pulmonary Healing Signaling Pathway 2.21
Leukotriene Biosynthesis 2.18
Phototransduction Pathway 2
Role of MAPK Signaling in Promoting the Pathogenesis of Influenza 1.85
Stearate Biosynthesis I (Animals) 1.82
Bladder Cancer Signaling 1.81
p38 MAPK Signaling 1.78
Phospholipases 1.75
Estrogen Receptor Signaling 1.71
Leukocyte Extravasation Signaling 1.66
Methylglyoxal Degradation III 1.63
Coenzyme A Biosynthesis 1.61
Colorectal Cancer Metastasis Signaling 1.6
Axonal Guidance Signaling 1.6
Ubiquinol-10 Biosynthesis (Eukaryotic) 1.6
CCR3 Signaling in Eosinophils 1.59
Gustation Pathway 1.58
Maturity Onset Diabetes of Young (MODY) Signaling 1.58
Opioid Signaling Pathway 1.56
Phagosome Formation 1.55
HIF1α Signaling 1.54
IL-8 Signaling 1.51
G-Protein Coupled Receptor Signaling 1.51
VEGF Family Ligand-Receptor Interactions 1.49
Agranulocyte Adhesion and Diapedesis 1.49
ERK/MAPK Signaling 1.49
Corticotropin Releasing Hormone Signaling 1.46
Endocannabinoid Neuronal Synapse Pathway 1.46
Interferon Signaling 1.45
Antigen Presentation Pathway 1.39
eNOS Signaling 1.37
Osteoarthritis Pathway 1.35
HOTAIR Regulatory Pathway 1.34
NRF2-mediated Oxidative Stress Response 1.33
Aldosterone Signaling in Epithelial Cells 1.33
SPINK1 Pancreatic Cancer Pathway 4.38
Retinol Biosynthesis 2.49
Acute Phase Response Signaling 2.16
Fatty Acid α-oxidation 2.1
Growth Hormone Signaling 1.97
Pregnenolone Biosynthesis 1.89
Histidine Degradation VI 1.86
Coagulation System 1.68
Oxidized GTP and dGTP Detoxification 1.6
Thyroid Hormone Metabolism II (via Conjugation and/or Degradation) 1.57
B Cell Activating Factor Signaling 1.51
IL-23 Signaling Pathway 1.45
Dermatan Sulfate Biosynthesis (Late Stages) 1.44
MSP-RON Signaling In Macrophages Pathway 1.4
Chondroitin Sulfate Biosynthesis (Late Stages) 1.39
FXR/RXR Activation 1.32
IL-6 Signaling 1.31
24h
72h
-log(p-value)
5
4
3
2

**Figure S8. Development and cancer pathways impacted by *her3* knockout.** Full list of pathways significantly enriched with differentially expressed genes from her3_38aa. -log(p-values) are displayed for each pathway per timepoint.
